# Evaluation of an image-derived input function for kinetic modeling of nicotinic acetylcholine receptor-binding PET ligands

**DOI:** 10.1101/2022.08.05.502975

**Authors:** Matthew Zammit, Chien-Min Kao, Hannah J. Zhang, Nathanial Holderman, Samuel Mitchell, Eve Tanios, Vincent Zhang, Mohammed Bhuiyan, Richard Freifelder, William N. Green, Jogeshwar Mukherjee, Chin-Tu Chen

## Abstract

Development of positron emission tomography (PET) radiotracers that bind with high-affinity to α4β2-type nicotinic receptors (α4β2Rs) allows for *in vivo* investigations of the mechanisms underlying nicotine addiction and smoking cessation. One challenge associated with preclinical PET imaging involves the lack of true tissue reference regions free of specific tracer binding in the rodent brain, impeding accurate quantification of the tracer binding potential. Here, we investigate the use of an image-derived arterial input function for kinetic analysis of radiotracer binding in male and female mice. Two radiotracers were explored in this study: 2-[^18^F]FA85380 (2-FA), which displays similar pKa and binding affinity to the smoking cessation drug varenicline (Chantix), and [^18^F]Nifene, which displays similar pKa and binding affinity to nicotine. For both radiotracers, time-activity curves of the left ventricle of the heart displayed similar standardized uptake values (SUVs) across wild type mice, mice lacking the β2 subunit for tracer binding, and acute nicotine-treated mice, whereas typical reference tissue SUVs displayed high variation between groups. Binding potential values estimated from a two-tissue compartment model (2TCM) fit of the data with the image-derived input function were significantly higher than estimates from reference tissue-based estimations. Rate constants of radiotracer dissociation were very slow for 2-FA and very fast for Nifene, similar to the *in vitro* dissociation rates reported for varenicline and nicotine, respectively. We conclude that use of an image-derived input function for kinetic modeling of nicotinic PET ligands improves quantification compared to reference tissue-based methods, and that the chemical properties of 2-FA and Nifene are suitable to study receptor response to nicotine addiction and smoking cessation therapies.

## Introduction

Tobacco use is the leading cause of preventable deaths in the United States and one of the prominent causes of nicotine addiction (SurgeonGeneral, 2014). Nicotine permeates the blood brain barrier and binds to high-affinity nicotinic acetylcholine receptors (nAChRs) containing α4 and β2 subunits (α4β2Rs) (Albuquerque et al., 2009). Chronic exposure to nicotine causes upregulation of α4β2Rs, in which increases in both the density of high-affinity binding sites and the functional response of α4β2Rs are observed (Benwell et al., 1988; Breese et al., 1997; Marks et al., 1983; Schwartz and Kellar, 1983). In addition, the process of nicotine-induced α4β2R upregulation has been linked to nicotine addiction (Govind et al., 2012, 2009; Lewis and Picciotto, 2013; Vezina et al., 2007).

Nicotine, and other weak-base ligands of α4β2Rs, such as the smoking cessation drug varenicline (Chantix), rapidly reach equilibrium in intracellular organelles and concentrate in acidic organelles (Brown and Garthwaite, 1979; Govind et al., 2017). The high pKa and binding affinity of varenicline causes selective trapping of this ligand inside intracellular acidic vesicles containing high-affinity α4β2Rs (Govind et al., 2017). Alternatively, nicotine concentrates inside these acidic vesicles but does not become trapped due to its lower pKa and lower binding affinity, and is rapidly released from the vesicles (Govind et al., 2017). Nicotine-induced upregulation increases the number of high-affinity binding sites within the acidic vesicles and increases the number of acidic vesicles, allowing for the vesicles to trap varenicline in higher concentrations. Recently, our *in vitro* studies suggest these acidic vesicles to be Golgi satellites (GSats), a novel intracellular compartment in neurons and neuronal dendrites that contain high α4β2R density, which increase in number following exposure to nicotine (Govind et al., 2021; Zhang et al., 2022). While nicotine and varenicline bind to the same α4β2Rs, the residence time of nicotine in the brain is 1-2 hours, compared to the 4-5 day residence time of varenicline (Govind et al., 2017), which may be attributed to varenicline trapping inside GSats. It has been shown that dissipating the pH gradient across GSats with chloroquine diphosphate or ammonium chloride prevents trapping of varenicline in GSats, and under these conditions, exposure to varenicline results in similar α4β2R upregulation as observed with nicotine (Zhang et al., 2022). Cell pretreatment with chloroquine diphosphate or ammonium chloride did not affect the extent of α4β2R upregulation induced by nicotine exposure.

Of interest is to study the effects of nicotine addiction and smoking cessation *in vivo*. This can be achieved using positron emission tomography (PET), in which the binding of nicotinic ligands to α4β2Rs can be observed noninvasively through injection of nanomolar concentrations of radiolabeled nicotine analogs (Mukherjee et al., 2018). Initial PET studies imaged [^11^C]nicotine, however this ligand suffered from rapid dissociation of the receptor-ligand complex, high levels of nonspecific binding, and its accumulation in the brain was highly dependent on cerebral blood flow (Lundqvist et al., 1998; Mazière and Delforge, 1995; Sihver et al., 2000). The radiotracer 2-[^18^F]FA85380 (2-FA) was developed as a less toxic analog of epibatidine that overcomes the shortcomings of [^11^C]nicotine and binds with high affinity to α4β2Rs (Horti et al., 1996), but requires a prolonged imaging session to achieve accurate quantification (Chefer et al., 2003). [^18^F]Nifene was then developed as a ligand with moderate affinity to α4β2Rs to improve upon the slow kinetics of 2-FA (Pichika et al., 2006). Because the slow kinetics of 2-FA closely resemble epibatidine and varenicline, and the fast kinetics of Nifene resemble nicotine, these ligands can be used to monitor the mechanisms of smoking cessation and nicotine addiction *in vivo*. Our previous *in vitro* findings with these ligands found that chronic exposure to Nifene resulted in similar α4β2R upregulation as observed with nicotine, whereas exposure to 2-FA did not cause upregulation, similar to varenicline (Zhang et al., 2022). The high pKa and high affinity of 2-FA likely result in the same GSat trapping observed with varenicline, and pH dissipation across GSats with chloroquine diphosphate or ammonium chloride prevented 2-FA trapping and caused significant α4β2R upregulation following exposure to 2-FA. This phenomenon was also observed *in vivo* using PET, in which mice pretreated with chloroquine diphosphate showed reduced binding of 2-FA, while Nifene binding was unaffected (Zhang et al., 2022).

One challenge associated with PET quantification of 2-FA and Nifene is the lack of a suitable reference tissue due to the abundance of α4β2Rs in the brain. Typically, the cerebellum is chosen as a reference tissue due to its low uptake and rapid washout of α4β2R-binding radiotracers. In nonhuman primates, the cerebellum was validated as a suitable reference tissue for α4β2R-binding radiotracers (Hillmer et al., 2011), however, in rodents, radiotracer concentrations in the cerebellum can be displaced by nicotine or lobeline injection (Hillmer et al., 2013; Vaupel et al., 2007), indicative of some specific binding signal. Human imaging studies incorporated the corpus collosum as a reference tissue, however this needs to be further validated using a nicotine challenge to measure the nicotine displaceable component (Mukherjee et al., 2018). It is speculated that the corpus collosum may be a suitable reference tissue free of specific binding in rodents (Kant et al., 2011), however, partial volume effects may negatively affect quantification in this region due to its small volume and the low spatial resolution of PET. Because of the lack of a true tissue reference region for rodent imaging, studies incorporating the cerebellum as a reference tissue likely underestimate the true binding potential. Apart from performing arterial cannulation to measure the input function, use of an image-derived input function for kinetic radiotracer analysis is speculated as a noninvasive alternative to improve image quantification (Zanotti-Fregonara et al., 2011). Studies incorporating image-derived input functions have shown that the time-activity curve (TAC) from the left ventricle of the heart was an accurate representation of arterial blood, obviating the need for arterial cannulation (Ferl et al., 2007; Tantawy and Peterson, 2010).

Here, we explore use of the left ventricle as an image-derived input function for quantification of 2-FA and Nifene PET images of mice. Left ventricle TACs were compared between wild type, β2-knockout, and acute nicotine-treated mice to explore how different mouse models influence radiotracer activity concentration in the blood pool. Using a two-tissue compartment model fit (2TCM) of the PET data, rate constants of radiotracer association and dissociation were calculated to determine the radiotracer binding potential. Binding potential values were then directly compared against estimates derived using the cerebellum reference tissue previously explored for these radiotracers. Finally, 2TCM fit simulations were performed to assess the stability of the rate constant estimates using the left ventricle TAC input function.

## Methods

### Animals

For 2-FA PET, 5 male and 5 female wild type mice (WT), 8 male and 3 female β2 nAChR knockout (KO) mice, and 3 male and 3 female acute nicotine-treated (AN) mice were imaged. For Nifene PET, 7 male and 5 female WT mice, 5 male and 2 female KO mice, and 2 male and 2 female AN mice were imaged. The male and female KO mice and their WT littermates were generated in house by breeding a heterozygous pair on the C57BL/6J background purchased from the Jackson Lab (Bar Harbor, ME) (Picciotto et al., 1995). Additional male and female WT mice at the same age were purchased directly from the Jackson lab and used in the same manner as the WT littermates. Animals were housed in The University of Chicago Animal Research Resources Center. The Institutional Animal Care and Use Committee of the University of Chicago, in accordance with National Institutes of Health guidelines, approved all animal procedures. Mice were maintained at 22–24°C on a 12:12-h light-dark cycle and provided food (standard mouse chow) and water ad libitum. All mice were 3-10 months of age. The number of the animals imaged, and their body weights are summarized in Table 1.

**Table 1.** Information on the animals imaged with 2-FA and Nifene.

|  | **WT** | **KO** | **AN** |
| --- | --- | --- | --- |
| **# Imaged with 2-FA** | 10 (5M/5F) | 11 (8M/3F) | 6 (3M/3F) |
| **# Imaged with Nifene** | 12 (7M/5F) | 7 (5M/2F) | 4 (2M/2F) |
| **Male body weight (g)** | 30.5 ± 0.6 | 30.6 ± 0.8 | 27.2 ± 2.2 |
| **Female body weight (g)** | 21.6 ± 0.7 | 21.0 ± 0.5 | 18.8 ± 0.8 |

### Radiotracer Syntheses

Syntheses of both [^18^F]2-FA and [^18^F]Nifene were carried out at the Cyclotron Facility of The University of Chicago. 2-FA was synthesized from the commercially available precursor, 2-TMA-A85380. Nifene was synthesized from the precursor N-BOC-nitroNifene. An IBA Synthera V2 automatic synthesis module equipped with Synthera preparative HPLC was used for the radiolabeling and purification inside a Comecer hot cell. The radiochemical yield of 2-FA was 34% (decay corrected) with specific activities >3,000 mCi/μmole and radiochemical purity > 99%. The radiochemical yield of Nifene was 6.3% (decay corrected) with specific activities >3,000 mCi/μmole and radiochemical purity > 99%.

### PET/CT Imaging

The imaging protocols were designed based on previous reports for 2-FA and Nifene (Constantinescu et al., 2013; Horti et al., 2000; Pichika et al., 2006) and our preliminary experiments fine-tuning the protocol (data not shown). An intraperitoneal (IP) catheter was placed at the lower right abdominal area of each mouse before imaging. The animal was then placed into the β-Cube preclinical microPET imaging system (Molecubes, Gent, Belgium) in a small animal holder. The tracer was delivered in 200 μL isotonic saline via the IP catheter and followed with addition of 100 μL of fresh saline. The injected doses are summarized in Table 1. For AN mice, 0.5 mg/kg body weight of nicotine was injected IP 15 minutes before radiotracer injection. Whole body imaging was acquired with a 133 mm x 72 mm field of view (FOV) and an average spatial resolution of 1.1 mm at the center of the FOV (Krishnamoorthy et al., 2018). List-mode data was recorded for 180 minutes for 2-FA and 60 minutes for Nifene followed by a reference CT image on the X-Cube preclinical microCT imaging system (Molecubes, Gent, Belgium). The images were reconstructed using an OSEM reconstruction algorithm that corrected for attenuation, randoms and scatter with an isotropic voxel size of 400 μm. The re-binned frame rate for 2-FA was 10×60s-17×600s and the frame rate for Nifene was 12×10s-18×60s-8×300s. CT images were reconstructed with a 200 μm isotropic voxel size and used for anatomic co-registration, scatter correction and attenuation correction. Animals were maintained under 1-2% isoflurane anesthesia in oxygen during imaging. Respiration and temperature were constantly monitored and maintained using the Molecubes monitoring interface, and a Small Animal Instruments (SAII Inc, Stoney Brook, NY) set up. All animals survived the imaging session.

### Image quantification

For each mouse imaged with 2-FA or Nifene, all PET frames were averaged and coregistered with the anatomical CT using VivoQuant (Invicro, Boston, MA). The resulting transformations were then applied to each individual PET image frame to align the PET with the CT. Brain regional analysis was performed using a 3-dimensional mouse brain atlas available through VivoQuant which is based on the Paxinos-Franklin atlas registered to a series of high-resolution magnetic resonance images with 100 μm near isotropic data that has been applied in other studies (Hesterman et al., 2019; Mazur et al., 2019; Slavine et al., 2017). The brain atlas was warped into the CT image space and used for volume of interest (VOI) extraction of the whole brain, cerebellum, thalamus and midbrain from the PET images. Regional radioactivity concentrations were converted into standardized uptake values (SUVs) by normalizing the signal by the injected dose of the radiotracer and the body weight of the animal. In addition, a spherical VOI was hand-drawn over the left ventricle of the heart to act as an image-derived input function for the kinetic modeling analyses.

### Radiotracer kinetic modeling

The two-tissue compartmental model (2TCM) is described by the following 1st order ordinary differential equations:

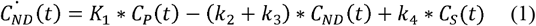

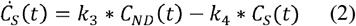

where C_ND_(t) and C_S_(t) are the concentration of the PET tracer in the non-displaceable and specifically bound compartments, respectively, and C_P_(t) is the concentration of PET tracer in the plasma compartment. K_1_, k_2_, k_3_, and k_4_ are the rate constants of PET tracer transport between the compartments. Using radioactivity time-activity curves (TACs) for all mice with the left ventricle TAC as an input function, the 2TCM was fit to the data to solve for the rate constants using an in-house Python software. Details regarding the in-house Python software are presented in the Appendix. To test the validity of the Python software, the 2TCM was also applied to the data using the built-in solver provided through the π.PMOD software. Taking the ratio of k_3_ and k_4_ resulted in an estimate of the binding potential (BP_ND_) which was determined for the thalamus, midbrain and cerebellum. Previous imaging studies of 2-FA and Nifene utilized the distribution volume ratio (DVR) as the outcome measure of specific binding, which is related to BP_ND_ via the following formula:

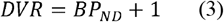

For the current study, the DVR was determined for the thalamus and midbrain using Logan graphical analysis (Logan et al., 1996) with the cerebellum acting as a tissue reference region. Using the concentration TACs generated from the VOIs from the VivoQuant brain atlas, a simplified reference tissue model (SRTM) (Lammertsma and Hume, 1996) was used to compute the *k*_2_^′^value for each region. Then a Logan plot was created, and the slope used as the DVR.

### 2TCM simulations

To validate the repeatability of the in-house Python solver for the 2TCM, the 2TCM was solved for a series of simulated 2-FA and Nifene data. For both 2-FA and Nifene, independently, regional TACs were taken from a single WT, KO and AN mouse. For each time point of the TAC, artificial noise was applied to the radioactivity concentration value with a noise level of 0.2 (noise standard deviation relative to the amplitude) such that the noise level was proportional to the amplitude/time duration. The 2TCM fit was then applied to the noise-induced thalamus and cerebellum TACs using the noise-induced left ventricle TAC as the input function. This process was repeated over 50 iterations for each mouse and radiotracer. Values for K_1_, k_2_, k_3_, k_4_ and BP_ND_ were then compared against the true values derived for each mouse.

### Statistical analyses

All statistical analyses were performed using R v3.0 (The R Project for Statistical Computing). Peak SUVs (max SUV across entire scan duration) and the mean SUVs from the final 30-minute image frames of the left ventricle were compared between WT, KO and AN mice using analysis of variance (ANOVA). For 2-FA and Nifene independently, K_1_ values from the thalamus and midbrain were compared between WT, KO and AN mice using ANOVA with post hoc Tukey’s HSD test for multiple comparisons. The ANOVA and post hoc tests were then repeated for k_2_, k_3_, k_4_, and BP_ND_ values. For both radiotracers, the 2TCM rate constants from the thalamus and midbrain determined from the in-house Python solver were directly compared to the rate constants determined from PMOD for the WT, KO and AN mice, independently using paired samples t-tests. BP_ND_+1 values determined from the 2TCM fit (left ventricle input function) were then directly compared with DVR values from Logan analysis (cerebellum reference tissue) using paired samples t-tests.

## Results

### Left ventricle comparisons

2-FA and Nifene time-activity curves (TACs) of the left ventricle, thalamus, midbrain and cerebellum for all mice are provided in Figure 1. For 2-FA, peak SUVs in the left ventricle (presented as mean(SEM)) for WT mice (0.55(0.055)), KO mice (0.47(0.065)) and AN mice (0.569(0.051)) showed no significant differences (ANOVA F(df): 0.62(2); p-value: 0.54). Mean SUVs from the final 30-minute data frames of WT mice (0.077(0.0091)), KO mice (0.081(0.015)) and AN mice (0.074(0.0059)) showed no significant differences (ANOVA F(df): 0.75(2); p-value: 0.93). For Nifene, peak SUVs in the left ventricle (presented as mean(SEM)) for WT mice (0.98(0.061)), KO mice (1.4(0.34)) and AN mice (1.2(0.073)) showed no significant differences (ANOVA F(df): 1.38(2); p-value: 0.27). Mean SUVs from the final 30-minute data frames of WT mice (0.41(0.041)), KO mice (0.45(0.048)) and AN mice (0.50(0.058)) showed no significant differences (ANOVA F(df): 0.64(2); p-value: 0.54).

**Figure 1.**
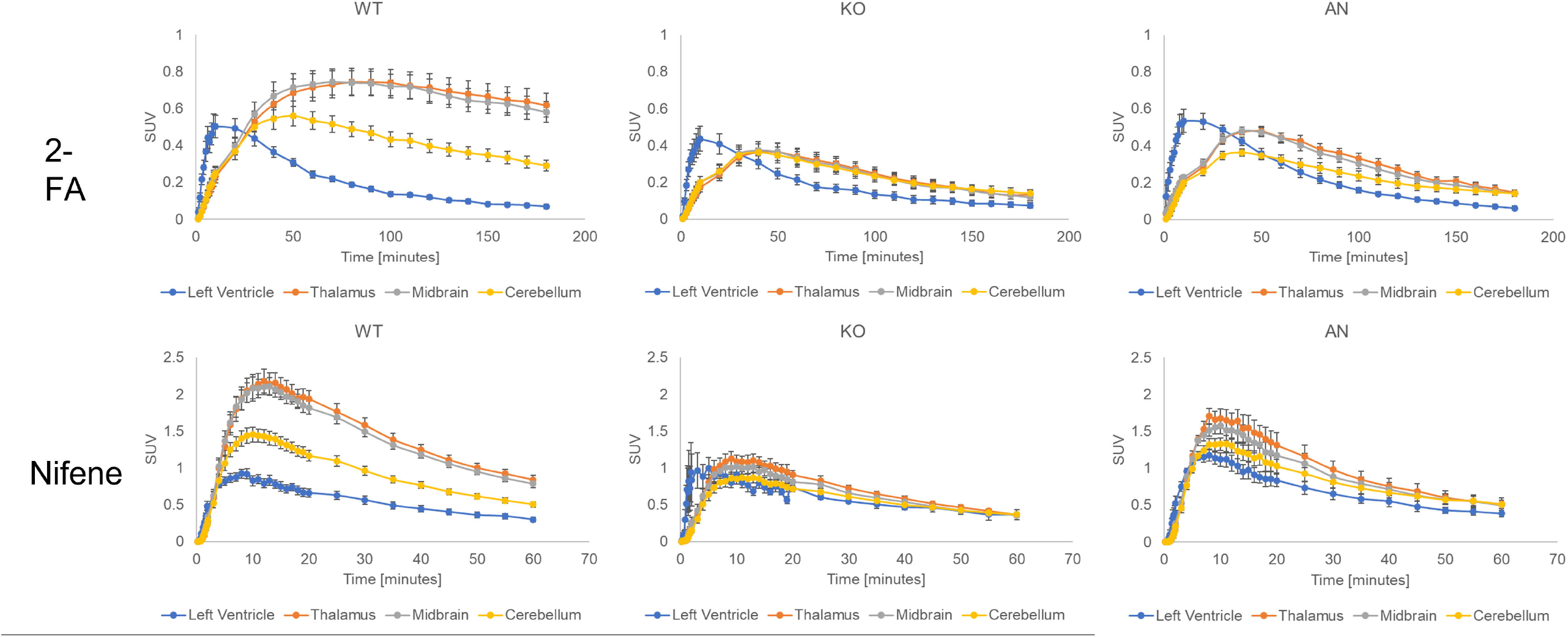
2-FA and Nifene standardized uptake value (SUV) time-activity curves (TACs) for WT, KO and AN mice. Error bars represent the standard error of the mean.

### Binding potential comparisons

Using the left ventricle TAC as an input to the 2TCM, BP_ND_ values were calculated for the thalamus, midbrain and cerebellum of WT, KO and AN mice by taking the ratio of k_3_ and k_4_ (Figure 2). From ANOVA with post hoc Tukey’s HSD, 2-FA BP_ND_ values were significantly higher in the thalamus, midbrain and cerebellum of WT mice compared to KO and AN mice (p<0.05), while no significant difference was observed between KO and AN mice (p>0.05). For Nifene, BP_ND_ values were significantly higher in the thalamus, midbrain and cerebellum of WT mice compared to KO and AN mice (p<0.05), while no significant difference was observed between KO and AN mice (p>0.05).

**Figure 2.**
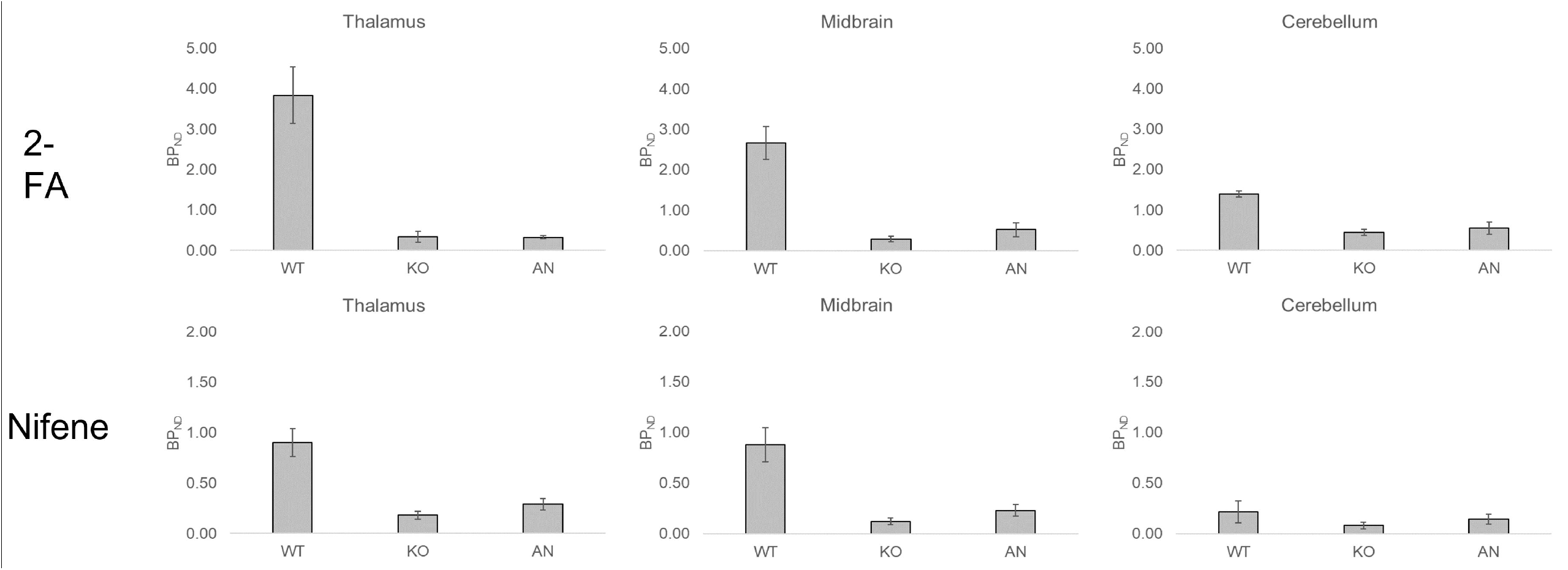
2-FA and Nifene BP_ND_ values for the thalamus, midbrain and cerebellum of WT, KO and AN mice.

### 2TCM rate constants

Table 2 provides the rate constant estimates with ANOVA F-statistics from the 2TCM for the thalamus and midbrain imaged with 2-FA and Nifene. For 2-FA, significant differences were observed between WT, KO and AN mice for K_1_ values in the thalamus, but not the midbrain. No significant differences were observed with k_2_ values in both the thalamus and midbrain across all groups. Significant differences between groups for both regions were observed for k_3_ and k_4_ values. Post hoc Tukey’s HSD revealed that the WT mice had higher estimates of k_3_ and lower estimates of k_4_ compared to KO and AN mice. In addition, Tukey’s HSD revealed no significant difference in k_3_ and k_4_ values between KO and AN mice. For Nifene, significant differences were observed between mice for K_1_ values in the thalamus and midbrain. From Tukey’s HSD, the significance was driven by the comparison between WT and KO mice. KO and AN mice, and WT and AN mice showed no significant difference between K_1_ estimates. No significant difference in k_2_ values were observed between mice in the thalamus, but midbrain k_2_ values were significantly different. For the midbrain, k_2_ values in WT mice were significantly higher than in the KO mice, and no differences were observed between the other pairings of mice. Significant difference in k_3_ values were observed between mice in the thalamus and midbrain. For both the thalamus and midbrain, no significant differences were observed for k_4_ values between mice.

**Table 2.** Rate constants for WT, KO and AN mice imaged with 2-FA and Nifene.

| **Radiotracer** | **Region** | **Rate constant (1/min)** | **WT** | **KO** | **AN** | **ANOVA F(df)** | **p-value** |
| --- | --- | --- | --- | --- | --- | --- | --- |
| **2-FA** | **Thalamus** | K<sub>1</sub> | 0.098(0.013) | 0.066(0.0061) | 0.060(0.0036) | **3.88(2)** | **0.035** |
|  |  | k<sub>2</sub> | 0.075(0.015) | 0.055(0.0066) | 0.050(0.0046) | 1.37(2) | 0.27 |
|  |  | k<sub>3</sub> | 0.029(0.0064) | 0.0049(0.0016) | 0.0054(0.00080) | **9.75(2)** | **0.00080** |
|  |  | k<sub>4</sub> | 0.0077(0.0013) | 0.020(0.0037) | 0.016(0.0017) | **5.48(2)** | **0.011** |
|  | **Midbrain** | K<sub>1</sub> | 0.11(0.017) | 0.076(0.0087) | 0.062(0.0051) | 2.67(2) | 0.090 |
|  |  | k<sub>2</sub> | 0.076(0.020) | 0.066(0.0087) | 0.051(0.0055) | 0.54(2) | 0.59 |
|  |  | k<sub>3</sub> | 0.024(0.0062) | 0.0060(0.0012) | 0.0031(0.00070) | **6.70(2)** | **0.0049** |
|  |  | k<sub>4</sub> | 0.0084(0.0016) | 0.026(0.0037) | 0.0093(0.0022) | **11.70(2)** | **0.00029** |
| **Nifene** | **Thalamus** | K<sub>1</sub> | 1.66(0.24) | 0.55(0.089) | 1.25(0.072) | **7.82(2)** | **0.0027** |
|  |  | k<sub>2</sub> | 1.23(0.26) | 0.51(0.074) | 1.09(0.068) | 3.05(2) | 0.068 |
|  |  | k<sub>3</sub> | 0.43(0.11) | 0.047(0.011) | 0.14(0.024) | **4.96(2)** | **0.017** |
|  |  | k<sub>4</sub> | 0.50(0.12) | 0.27(0.034) | 0.51(0.12) | 1.71(2) | 0.20 |
|  | **Midbrain** | K<sub>1</sub> | 1.94(0.31) | 0.50(0.080) | 1.28(0.099) | **8.00(2)** | **0.0024** |
|  |  | k<sub>2</sub> | 1.52(0.35) | 0.47(0.069) | 1.14(0.061) | **3.54(2)** | **0.047** |
|  |  | k<sub>3</sub> | 0.50(0.13) | 0.029(0.0085) | 0.12(0.026) | **5.75(2)** | **0.0098** |
|  |  | k<sub>4</sub> | 0.60(0.14) | 0.26(0.046) | 0.60(0.077) | 2.21(2) | 0.13 |

### Rate constant comparisons between in-house Python solver and PMOD solver

Table 3 displays the 2-FA rate constant estimates from the Python and PMOD 2TCM solvers, and the pairwise t-test comparisons between them. For WT mice, there were no significant differences between estimates of K_1_, k_2_, k_3_ or k_4_ in the thalamus or midbrain between methods. PMOD estimates for K_1_, k_3_ and k_4_ were significantly higher than the Python solver in KO mice, and PMOD estimates for k_3_ and k_4_ were higher in AN mice. Because KO and AN mice display no specific binding of 2-FA, estimates of these rate constants display high variance, which may influence the trend of higher mean values. Table 4 displays the Nifene rate constant estimates from the Python and PMOD 2TCM solvers, and the pairwise t-test comparisons between them. For WT mice, there were no significant differences between estimates of k_3_ or k_4_ in the thalamus or midbrain between methods. However, K_1_ and k_2_ estimates from the Python solver were significantly higher than the PMOD estimates. KO and AN mice revealed no significant differences between rate constant values in the thalamus and midbrain. Lack of Nifene specific binding in KO and AN mice contributes to the higher variance in these measurements for the Python and PMOD solvers.

**Table 3.** 2-FA rate constants measured using the in-house Python solver and the PMOD solver compared using pairwise t-tests.

| Group | Rate constant (1/min) | Region | 2TCM Python | 2TCM PMOD | T-value | p-value |
| --- | --- | --- | --- | --- | --- | --- |
| WT | K<sub>1</sub> | Thalamus | 0.098(0.013) | 0.13(0.022) | 1.17 | 0.27 |
|  |  | Midbrain | 0.11(0.017) | 0.12(0.014) | 1.23 | 0.25 |
|  | k<sub>2</sub> | Thalamus | 0.075(0.015) | 0.19(0.085) | 1.24 | 0.24 |
|  |  | Midbrain | 0.076(0.020) | 0.13(0.028) | 1.61 | 0.14 |
|  | k<sub>3</sub> | Thalamus | 0.029(0.0064) | 0.071(0.021) | 1.69 | 0.12 |
|  |  | Midbrain | 0.024(0.0062) | 0.043(0.0076) | 2.26 | 0.051 |
|  | k<sub>4</sub> | Thalamus | 0.0077(0.0013) | 0.026(0.017) | 0.98 | 0.35 |
|  |  | Midbrain | 0.0084(0.0016) | 0.013(0.0032) | 1.28 | 0.23 |
| KO | K<sub>1</sub> | Thalamus | 0.066(0.0061) | 0.14(0.030) | **2.39** | **0.038** |
|  |  | Midbrain | 0.076(0.0087) | 0.18(0.041) | **2.46** | **0.034** |
|  | k<sub>2</sub> | Thalamus | 0.055(0.0066) | 0.44(0.18) | 1.97 | 0.077 |
|  |  | Midbrain | 0.066(0.0087) | 0.54(0.20) | 2.18 | 0.054 |
|  | k<sub>3</sub> | Thalamus | 0.0049(0.0016) | 0.090(0.023) | **3.49** | **0.0059** |
|  |  | Midbrain | 0.0060(0.0012) | 0.092(0.027) | **2.98** | **0.014** |
|  | k<sub>4</sub> | Thalamus | 0.020(0.0037) | 0.049(0.0060) | **5.39** | **0.00030** |
|  |  | Midbrain | 0.026(0.0037) | 0.044(0.0080) | **3.02** | **0.013** |
| AN | K<sub>1</sub> | Thalamus | 0.060(0.0036) | 0.34(0.13) | 1.85 | 0.12 |
|  |  | Midbrain | 0.062(0.0051) | 0.37(0.12) | 2.26 | 0.073 |
|  | k<sub>2</sub> | Thalamus | 0.050(0.0046) | 1.23(0.63) | 1.71 | 0.15 |
|  |  | Midbrain | 0.051(0.0055) | 1.19(0.47) | 2.20 | 0.079 |
|  | k<sub>3</sub> | Thalamus | 0.0054(0.00080) | 0.087(0.023) | **2.91** | **0.033** |
|  |  | Midbrain | 0.0031(0.00070) | 0.085(0.028) | **2.67** | **0.045** |
|  | k<sub>4</sub> | Thalamus | 0.016(0.0017) | 0.030(0.0028) | **3.14** | **0.026** |
|  |  | Midbrain | 0.0093(0.0022) | 0.031(0.0024) | **6.03** | **0.0018** |

**Table 4.** Nifene rate constants measured using the in-house Python solver and the PMOD solver compared using pairwise t-tests.

| Group | Rate constant (1/min) | Region | 2TCM Python | 2TCM PMOD | T-value | p-value |
| --- | --- | --- | --- | --- | --- | --- |
| WT | K<sub>1</sub> | Thalamus | 1.66(0.24) | 1.18(0.13) | **3.60** | **0.0042** |
|  |  | Midbrain | 1.94(0.31) | 1.45(0.17) | **2.83** | **0.016** |
|  | k<sub>2</sub> | Thalamus | 1.23(0.26) | 0.61(0.12) | **3.37** | **0.0062** |
|  |  | Midbrain | 1.52(0.35) | 0.85(0.15) | **2.33** | **0.040** |
|  | k<sub>3</sub> | Thalamus | 0.43(0.11) | 0.76(0.51) | 0.72 | 0.49 |
|  |  | Midbrain | 0.50(0.13) | 0.36(0.12) | 0.68 | 0.51 |
|  | k<sub>4</sub> | Thalamus | 0.50(0.12) | 1.63(0.62) | 2.09 | 0.060 |
|  |  | Midbrain | 0.60(0.14) | 1.42(0.56) | 1.77 | 0.10 |
| KO | K<sub>1</sub> | Thalamus | 0.55(0.089) | 0.52(0.12) | 0.73 | 0.48 |
|  |  | Midbrain | 0.50(0.080) | 0.50(0.11) | 0.0096 | 0.99 |
|  | k<sub>2</sub> | Thalamus | 0.51(0.074) | 0.63(0.29) | 0.52 | 0.62 |
|  |  | Midbrain | 0.47(0.069) | 0.45(0.098) | 0.33 | 0.75 |
|  | k<sub>3</sub> | Thalamus | 0.047(0.011) | 0.075(0.037) | 0.62 | 0.55 |
|  |  | Midbrain | 0.029(0.0085) | 0.068(0.043) | 0.87 | 0.41 |
|  | k<sub>4</sub> | Thalamus | 0.27(0.034) | 2.64(1.04) | 2.16 | 0.062 |
|  |  | Midbrain | 0.26(0.046) | 0.65(0.35) | 1.01 | 0.34 |
| AN | K<sub>1</sub> | Thalamus | 1.25(0.072) | 1.00(0.050) | 1.92 | 0.15 |
|  |  | Midbrain | 1.28(0.099) | 1.25(0.083) | 0.20 | 0.86 |
|  | k<sub>2</sub> | Thalamus | 1.09(0.068) | 0.86(0.15) | 1.09 | 0.35 |
|  |  | Midbrain | 1.14(0.061) | 1.29(0.21) | 0.57 | 0.61 |
|  | k<sub>3</sub> | Thalamus | 0.14(0.024) | 0.65(0.39) | 1.19 | 0.32 |
|  |  | Midbrain | 0.12(0.026) | 1.37(1.04) | 1.02 | 0.38 |
|  | k<sub>4</sub> | Thalamus | 0.51(0.12) | 4.23(1.73) | 1.90 | 0.15 |
|  |  | Midbrain | 0.60(0.077) | 3.83(1.62) | 1.80 | 0.17 |

### 2TCM comparisons to Logan graphical analysis

The 2TCM estimate of BP_ND_+1 from the Python solver using a left ventricle input function was directly compared to the Logan graphical analysis estimate of DVR using a cerebellum reference tissue. Table 5 displays the pairwise comparisons between BP_ND_+1 and DVR. For 2-FA, BP_ND_+1 values were significantly higher than DVRs in the thalamus and midbrain of WT mice. KO and AN mice also displayed significantly higher BP_ND_+1 values compared to DVR, however the values are low and indicative of no specific binding signal. As illustrated in Figure 2, BP_ND_ in the cerebellum of WT mice was significantly higher than observed in KO and AN mice, indicative of some 2-FA specific binding in this region. This cerebellar specific binding signal results in an underestimation of the true binding potential in the thalamus and midbrain when estimated from Logan DVR. For Nifene, no significant difference between BP_ND_+1 and DVR were observed in the thalamus or midbrain of WT mice, likely because the levels of cerebellar specific binding are much lower compared to 2-FA as shown in Figure 2 and further illustrated by the SUV TACs in Figure 1. Nifene BP_ND_+1 in KO and AN mice also displayed higher values compared to DVR, however, these values are low and indicative of no radiotracer specific binding.

**Table 5.** Pairwise comparisons between BP_ND_+1 from the 2TCM and DVR from Logan graphical analysis for 2-FA and Nifene.

| **Radiotracer** | **Group** | **Region** | **$BP_{ND+1}$** | **DVR** | **T-value** | **p-value** |
| --- | --- | --- | --- | --- | --- | --- |
| **2-FA** | WT | Thalamus | 4.84(0.70) | 2.35(0.15) | **3.94** | **0.0034** |
|  |  | Midbrain | 3.66(0.41) | 1.96(0.080) | **4.15** | **0.0025** |
|  | KO | Thalamus | 1.34(0.14) | 0.97(0.019) | **2.57** | **0.028** |
|  |  | Midbrain | 1.29(0.064) | 0.97(0.011) | **4.78** | **0.00075** |
|  | AN | Thalamus | 1.33(0.037) | 1.15(0.051) | **2.73** | **0.042** |
|  |  | Midbrain | 1.52(0.17) | 1.05(0.023) | 2.40 | 0.062 |
| **Nifene** | WT | Thalamus | 1.90(0.14) | 1.79(0.079) | 0.65 | 0.53 |
|  |  | Midbrain | 1.88(0.17) | 1.54(0.092) | 1.82 | 0.095 |
|  | KO | Thalamus | 1.18(0.038) | 0.94(0.039) | **5.03** | **0.0010** |
|  |  | Midbrain | 1.12(0.034) | 0.98(0.015) | **3.11** | **0.015** |
|  | AN | Thalamus | 1.29(0.056) | 0.93(0.016) | **5.79** | **0.010** |
|  |  | Midbrain | 1.23(0.056) | 0.99(0.0049) | **3.57** | **0.037** |

### Simulations

Simulations were performed on the PET data to evaluate the stability of the output parameters of the 2TCM fit when solved using the in-house Python solver. Using noise-induced TACs of the left ventricle, cerebellum and thalamus, the 2TCM fit was applied to the data to solve for BP_ND_, K_1_, k_2_, k_3_ and k_4_ values for sample WT, KO and AN mice. Figure 3 displays the results of the simulated 2-FA data. For the WT and KO mice, the true values of BP_ND_, K_1_, k_2_, k_3_ and k_4_ all fell within the 25^th^ and 75^th^ percentile of the simulated estimates. For the AN mice, the true values of BP_ND_, K_1_, k_2_, k_3_ and k_4_ all fell within the 25^th^ and 75^th^ percentile of the simulated estimates, except for the BP_ND_ of the thalamus. Figure 4 displays the results of the simulated Nifene data. For BP_ND_ values, the true values fell within the 25^th^ and 75^th^ percentile of the simulated estimates for the WT, KO and AN mice. For K_1_ and k_2_ values, the 2TCM fit slightly underestimated the true values for all groups of mice. For k_3_ and k_4_ values, the true values fell within the 25^th^ and 75^th^ percentile of the simulated estimates for the KO and AN mice, while the WT mice simulations slightly overestimated the true values.

**Figure 3.**
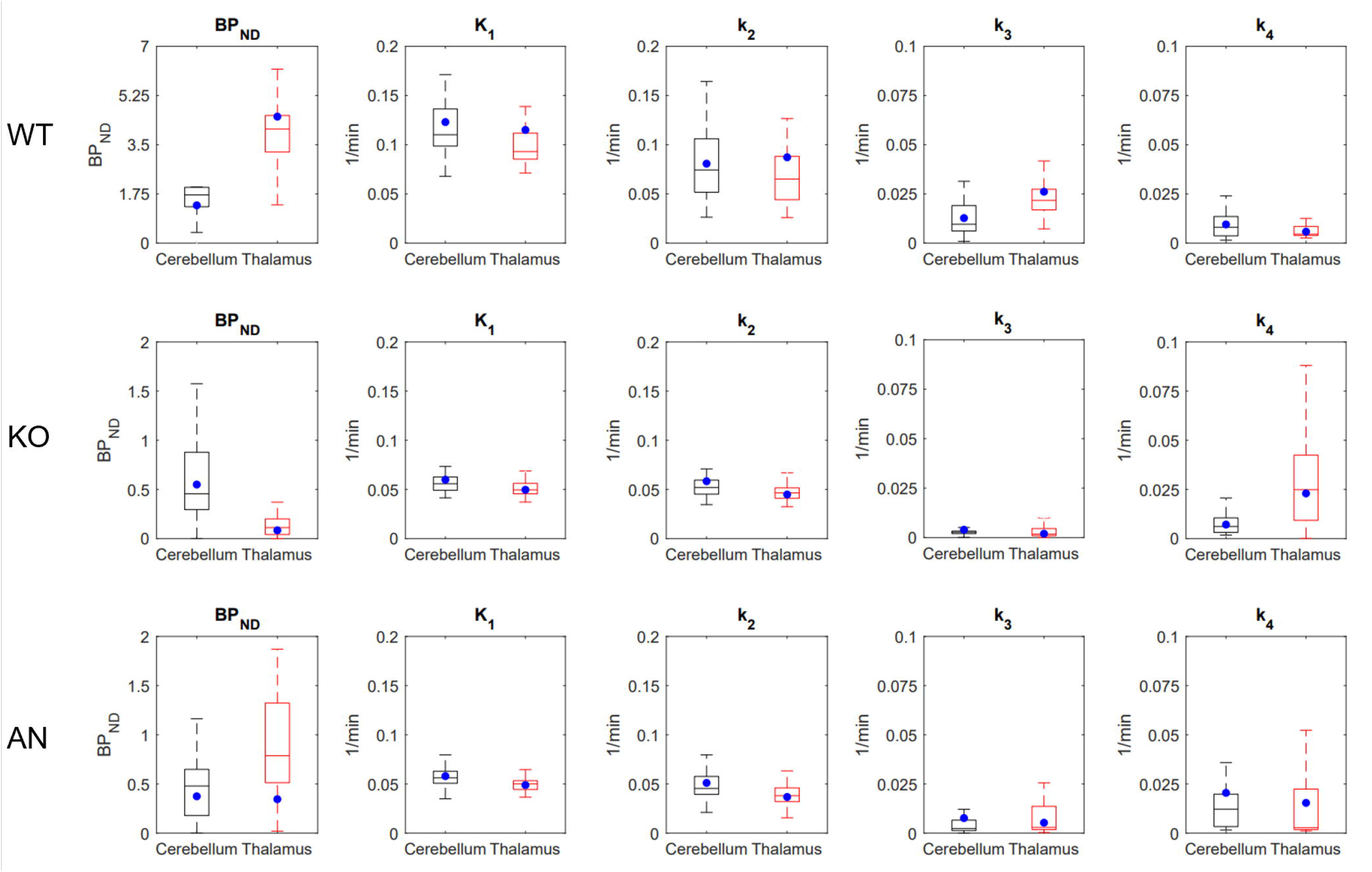
Distribution of simulated 2-FA BP_ND_, K_1_, k_2_, k_3_ and k_4_ estimates from WT, KO and AN mice in the cerebellum and thalamus using a left ventricle input function. The true values of each estimate are displayed as blue circles.

**Figure 4.**
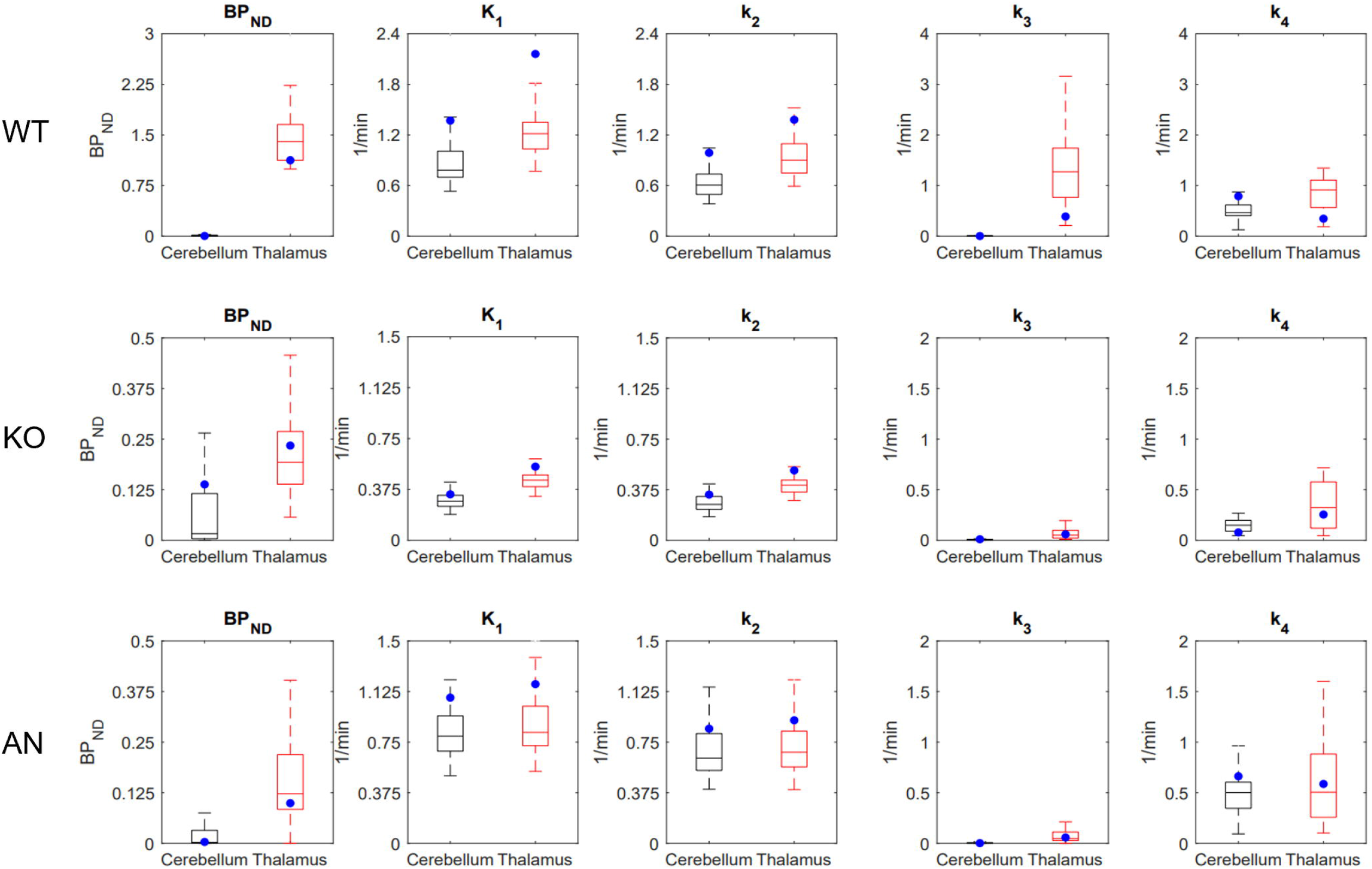
Distribution of simulated Nifene BP_ND_, K_1_, k_2_, k_3_ and k_4_ estimates from WT, KO and AN mice in the cerebellum and thalamus using a left ventricle input function. The true values of each estimate are displayed as blue circles.

## Discussion

This study is the first to explore the use of an image-derived input function for kinetic analysis of 2-FA and Nifene PET images in mice. An ROI of the left ventricle was chosen as the image-derived input function due to its accurate representation of the plasma fraction as shown in other studies (Ferl et al., 2007; Tantawy and Peterson, 2010; Zanotti-Fregonara et al., 2011). Across WT, KO and AN mice, no significant differences were observed between the SUV TACs of the left ventricle, suggesting this region as a suitable reference for these mouse models when imaging with 2-FA and Nifene. Previous studies using 2-FA and Nifene in rodents utilized the cerebellum as a tissue reference region and identified nicotine-displaceable signal in this region (Hillmer et al., 2013; Vaupel et al., 2007). The current study confirms the presence of α4β2Rs in the cerebellum, notably due to the higher binding potential values of 2-FA and Nifene in the WT group compared to the KO and AN groups. Thus, studies with rodents using the cerebellum as a tissue reference region will underestimate the true binding potential in target regions of interest, further emphasizing the need for improved quantification derived from arterial data. One limitation to our application of using an image-derived input function was delivery of the radiotracers through IP injection. IP injections result in a slower distribution of radioligand from the plasma compartment to target regions when compared to the more standard IV injection. As a result, 2TCM fitting of the radioactivity time course of Nifene, which has very rapid kinetics *in vivo*, was challenging as the data was better represented by a 1TCM. Compared to 2-FA, Nifene is more lipophilic and tends to concentrate in abdominal fat when injected IP, resulting in a slower plasma time course. Because the kinetics of 2-FA are very slow, use of an IP injection did not present a challenge in 2TCM fitting. Despite the challenges associated with IP injections, the 2TCM fit of the data using an image-derived input function resulted in more quantitatively accurate estimates of the binding potential for 2-FA and Nifene compared to tissue reference models. To further validate the use of a left ventricle image-derived input function, future studies incorporating arterial sampling through cannulation should be performed for comparison.

The current study incorporated use of an in-house Python solver and PMOD for the 2TCM fits of the PET data. The in-house Python solver was developed to derive more stable estimates of the rate constants for the KO and AN mouse groups. Due to the low binding levels of 2-FA and Nifene in these groups, rate constant estimates using PMOD had high uncertainties. Use of parameter optimization in the Python solver greatly reduced these uncertainties in the estimates, allowing for better comparisons to the rate constants derived for the WT mice. Importantly, no significant differences were observed between the Python solver and PMOD when fitting data from the WT mice, which displayed high levels of radiotracer binding. For the WT mice, estimates of k_3_ and k_4_ were large for Nifene, indicative of rapid binding and unbinding from α4β2Rs. Alternatively, 2-FA showed small estimates of k_3_ and very small estimates of k_4_, indicative of slow binding and very slow unbinding from α4β2Rs. This very slow unbinding of2-FA may partially be influenced by ligand trapping inside GSats, which has been shown previously *in vitro* and *in vivo* (Govind et al., 2021, 2017; Zhang et al., 2022). Of interest is to use these radiotracers to directly study the mechanisms of nicotine addiction and smoking cessation. *In vitro* cellular studies performed under similar *in vivo* conditions in mice found that the dissociation rate of nicotine to be 0.84 1/min (Lippiello et al., 1987), while the dissociation rate of epibatidine was 0.043 1/min (Whiteaker et al., 1998). The *in vivo* dissociation rate (k_4_) estimates of Nifene and 2-FA derived in the current study (Table 2) fall within the same order of magnitude as the *in vitro* estimates of nicotine and epibatidine, suggesting that these PET tracers may be useful to study addiction and smoking cessation mechanisms, especially those involving GSat trapping and release. Because 2-FA was only imaged for a 3-hour duration, a true estimate of 2-FA release from GSats could not be obtained, as the radiotracer did not reach equilibrium in the brain by the end of the scan. Previous *in vitro* work measured the dissociation of epibatidine over a 17-hour duration and found that there was a rapid component (unbinding from α4β2Rs) and a very slow component (release from GSats) (Govind et al., 2017). Because 2-FA is an analog of epibatidine, it is speculated that this slow component of dissociation is similar between ligands and can be measured *in vivo*. Future work should explore whether an estimate of 2-FA release from GSats can be derived, however, the radioactive decay of F-18 may present a challenge when imaging for long scan durations. Alternatively, future studies could incorporate single photon emission computed tomography (SPECT) scans using I-123 or I-125 labeled analogs of epibatidine to measure the release of ligand from GSats.

To directly compare to DVR estimates from tissue reference methods, BP_ND_+1 values from the 2TCM fit were compared to DVR from Logan graphical analysis (Logan et al., 1996) using a cerebellum reference tissue. For 2-FA, BP_ND_+1 was significantly higher than DVR for the thalamus and midbrain across all mouse groups. This difference is primarily due to the large BP_ND_ values present in the cerebellum, resulting in the Logan DVR underestimating the true binding potential. For Nifene, no significant difference between BP_ND_+1 and DVR in the thalamus and midbrain were observed for WT mice. This is likely because the levels of specific binding of Nifene in the cerebellum are much lower than observed with 2-FA (Figure 2). Despite the lack of significance, BP_ND_+1 values for the WT mice were higher than the observed DVR values, indicative of some levels of specific binding present in the cerebellum, resulting in DVR underestimating the true binding potential. BP_ND_+1 values for KO and AN mice were significantly higher than DVR values for both the thalamus and midbrain, however, these mice display very low binding of tracer.

To test the reliability of the Python solver for the 2TCM fitting, simulations were performed on the 2-FA and Nifene PET data. Using a mix of male and female mice from the WT, KO and AN groups, artificial Gaussian noise was applied to the TACs of the thalamus, cerebellum and left ventricle. The noise-induced TACs were then fit by the 2TCM and estimates of the rate constants and binding potential values were derived and compared to the true values derived from the noise-free data. For all groups of mice imaged with 2-FA, the true estimates of BP_ND_, K_1_, k_2_, k_3_ and k_4_ fell within the 25^th^ and 75^th^ percentiles of the simulated values. These findings suggest that the Python solver provides stable estimates of the radiotracer rate constants for 2-FA. For Nifene, the true estimates of BP_ND_ fell within the 25^th^ and 75^th^ percentiles of the simulated values for all groups of mice, however, estimates of K_1_ and k_2_ were underestimated and k_3_ and k_4_ were overestimated for the WT group only. Because the Nifene TACs were well approximated by the 1TCM, the 2TCM fit had higher levels of uncertainty affiliated with the outcome measures. This uncertainty coupled with the slow input function kinetics resulting from the IP injection likely resulted in the poor estimation of the rate constants for Nifene. Future work will evaluate the 2TCM fit with 2-FA and Nifene delivered via IV injection to improve the tracer kinetics.

In summary, we present evidence that use of an image-derived input function provides improved quantification of 2-FA and Nifene PET data compared to estimates from a tissue reference region. Consistent with the previous rodent work performed with these radiotracers, both 2-FA and Nifene display specific binding in the cerebellum, further emphasizing the need for analyses using an arterial input function to better estimate α4β2R density. Due to the similarity in structure and kinetics of 2-FA and Nifene to varenicline and nicotine, respectively, these radiotracers are ideal for studying the underlying mechanisms of nicotine addiction and smoking cessation *in vivo*.

## Appendix

Solving Eqs. (1) & (2) yields *C*_*t*_(*t*) = *h*(*t*) ⋆ *C*_*p*_(*t*) where the response function *h*(*t*) is given by (Gunn et al., 2001)

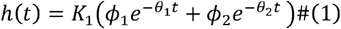

with

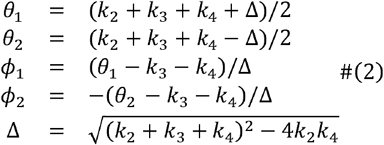

Therefore, *C*_*t*_(*t*) is determined by the rate constants and *C*_*p*_(*t*) It can be checked that 0 ≤ *ϕ*_1_, *ϕ*_2_ ≤ 1 and *ϕ*_1_ + *ϕ*_2_ = 1. Given the TAC for a VOI, the rate constants and *ν*_*p*_ for this VOI are the weighted least-squares solution given by:

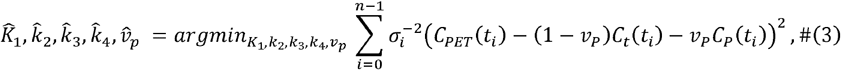

where *t*_*i*_ is the mid-time of the *i* th time frame and 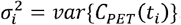. By setting *k*_3_ = *k*_4_ = 0, the above also yields solutions for the one-tissue compartmental model (1TCM) parameters *k*_1_, *k*_2_ and *ν*_*p*_

Within a scaling factor, the variance 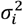 is estimated as follows. Let *y*_*ij*_ denote the PET intensity value before decay correction where *i* and *j* identify the image voxel and time frame respectively, and ⟨ *y*_*ij*_ ⟩ _*j:VOI*_ the average of *y*_*ij*_ for voxels in a VOI. Then, *C*_*PET*_(*t*_*i*_) = (*ca*_*i*_/ Δ*t*_*i*_) ⟨ *y*_*ij*_ ⟩_*j:VOI*_ where *C* is a factor for converting the image intensity to a quantity with a physical unit (e.g., radioactivity concentration), *a*_*i*_ is the correction factor for isotope decay, and Δ*t*_*i*_ is the duration of the frame. We assume *var* { *y*_*ij*_ } ∝ *E*{ *y*_*ij*_ }.Therefore,

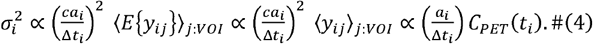

Let 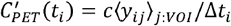 be the TAC before decay correction. Similarly, we obtain 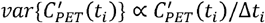. Therefore, when adding noise to a simulated noise-free TAC that includes isotope decay, the noise variance was proportional to the TAC value and inversely proportional to the frame duration, with the proportionality constant determined to yield 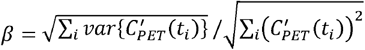 where *β* was the desired noise level given by the caller. 2TCM fitting was challenging for the KO and AN animal groups. On observing that, for them 1TCM could fit the tissue TACs reasonably, we examined and found that there are three situations when the 2TCM becomes approximately 1TCM and hence present ambiguities in the determining the solution. Assuming *k*_3_ + *k*_4_ ≪ *k*_2_ in Eqs. (1) and (2), we have

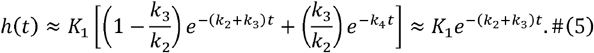

On the other hand, when *k*_3_ + *k*_4_ ≫ *k*_2_ we get

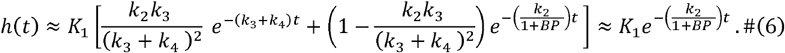

Both Eqs. (5) and (6) are a slightly perturbed exponential function. The third situation is when *θ*_1_ ∼ *θ*_2_ that occurs Δ ∼ 0 and requires *k*_3_ ∼ 0 and *k*_2_ ∼ *k*_4._ For the KO and AN groups, we expect *k*_3_ + *k*_4_ ≪ *k*_2_ and hence the solution given by Eq. (5). The fitting algorithm however could wrongly yield the solution Eq. (6) and obtain large *k*_3_ and *k*_4_ instead. To remove this ambiguity, we note that *ϕ*_1_ is close to 1 in Eq. (5), and close to 0 in Eq. (6). In addition, for both 2FA and Nifene we observe that *k*_2_ > *k*_3,_ *k*_4_ for the WT group, and one can show *ϕ*_1_ > 1/2 when *k*_2_ > *k*_3,_ *k*_4_. Therefore, the fitting algorithm accept the condition *ϕ*_*min*_ ≤ *ϕ*_1_ ≤ *ϕ*_*max*_ where 1/2 < *ϕ*_*min*_ < *ϕ*_*max*_ < 1 for rejecting the solutions with *k*_2_ ≪ *k*_3_ + *k*_4_. Specifically, we used 0.55 ≤ *ϕ*_1_ ≤ 0.95 for the WT animal group and 0.85 ≤ *ϕ*_1_ ≤ 0.95 for the KO and AN groups. However, this condition can not reject the *θ*_1_ *θ*_2_ solutions as their *ϕ*_1_ can assume any value. Therefore, as will be explained below, we will also seek to maximize Δ.

Another issue with our experimental data is related to the use of IP injection. Typically, IV injection is used and *C*_*P*_(*t*) shows a sharp peak within a few minutes post acquisition. With IP injection, in our data *C*_*P*_(*t*) showed a peak at about 10 and 20 minutes post acquisition with Nifene and 2FA respectively and decreased slowly. With a broader *C*_*P*_(*t*), generally a certain change in a rate constant or *ν*_*p*_ yields smaller changes in the tissue TACs. Consequently, the fitting result is more sensitive to data noise. The issue is even more challenging with weak binding as the approximate response function in Eq. (5) degenerate to depend on only three parameters *K*_1_ (*k*_3_/ *k*_2_), and *k*_4_. With IP injection, *C*_*P*_(*t*) also showed greater inter-subject variabilities. One way to help stabilize the fitting result is to define ranges for the fitting parameters. For this purpose, the in-house algorithm considers 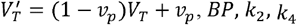 and *ν*_*p*_, where *V*_*T*_ = (*K*_1_/*k*_2_) (1 + *BP*) and *BP* = *k*_3_/*k*_4_, in place of *K*_1,_ *k*_2,_ *k*_3,_ *k*_4_ and *ν*_*p*_ because, as will be described below, we can define ranges for 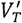 and *BP*.

The in-house algorithm began with a wide range *R*(*x*) for a parameter *x* that was unlikely to be violated. Specifically, 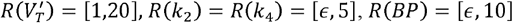, and *R*(*ν*_*p*_) = [*ε*,0.3] where ε is a small number. Subsequently, if another range*R*’(*x*) was specified for *x*, the range was updated to *R*(*x*) ∩ *R ′* (*x*). The following steps were taken to update the ranges. First, the slope *m* of the Logan plot, which is an estimate of 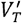, was obtained to define 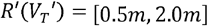. Next, the tissue TAC was fitted with the 1TCM model based on the parameters *K*_1,_ *k*_2,_ and *ν*_*p*._In this case, *R*(*K*_1_) = [*ε*,5] was employed and the updated 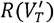 was applied to 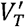 calculated from the fitting *K*_1,_ *k*_2_ and *ν*_*p*._ Using the resulting 1TCM estimates, 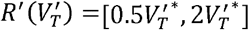 and 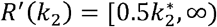, where 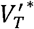 and 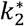 were the 1TCM solutions, were defined to update the fitting range for 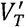 and *k*_2_. Subsequently, we performed 2TCM fitting for the cerebellum VOI that was known to have weak binding so that we could impose a small range for its *BP*. Specifically, we used *R*′ (*BP*) = [0,2]. The cerebellum result was then used to obtain *R*′ (*BP*) for other VOIs by employing the assumption commonly made in the reference-tissue methods: the non-displaceable distribution volume *V*_*ND*_ = *K*_1_/ *K*_2_ is identical for all VOIs. This assumption yielded 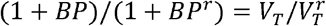, or 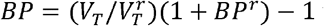 where 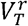 and *BP*^*r*^ are *V*_*T*_ and *BP* of the cerebellum. We therefore applied the range condition *R ′*(*BP*) = [0.5 *BP* ^*\**^,2.0 *BP* ^*\**^] where *BP* ^*\**^ is obtained for the VOI under examination using the above relationship. We also used 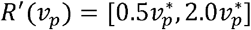, where 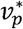 was the cerebellum value, by assuming all VOIs have similar *ν*_*p*_’s. We also set *V*_*ND*_ ≥ 1.

The resulting 2TCM fitting is a nonlinear optimization problem that incorporates range conditions on the optimization parameters and several other constraints. It was solved by employing the basinhopping algorithm provided in the Python scipy package with which the range conditions and constraints could be readily incorporated. As mentioned above, we need to avoid the *θ*_1_ ∼ *θ*_2_ solutions when the 2TCM is approximately 1TCM. This is achieved as follows. The basinhopping algorithm is a two-step global optimizer for functions that may have multiple local minima in their respective “basins”. The algorithm uses a local search algorithm to find the local minimum in a basin and random perturbations to jump basins to visit other local minima. We recorded all the local minima found by the algorithm and identified those whose weighted squared errors (WLS) are within 5% of the global minimal WLS. Among these potential solutions, the one having the maximum Δ, so that its *θ*_1_ and *θ*_2_ values are most distinct, was selected as the solution.

## Acknowledgements

This study was supported in part by the NIH grant R01 DA044760-01 to W.N.G., J.M., and C.C., RF1 AG029479 to J.M., and T32 DA043469 to M.Z. The authors acknowledge the assistance from the Integrative Small Animal Imaging Research Resources (iSAIRR) supported in part by the NIH grant P30 CA14500 and S10 OD025265, and from the Cyclotron Facility of the University of Chicago.

